# The sperm specific Na^+^,K^+^-ATPase α4 shows a highly structured and dynamic distribution at the sperm flagellum

**DOI:** 10.1101/2025.02.14.638303

**Authors:** Mumtarin J Oishee, Jeffrey P. McDermott, Gladis Sánchez, Gustavo Blanco

## Abstract

Na^+^,K^+^-ATPase α4 is a unique cell plasma membrane Na^+^ and K^+^ transporter of spermatozoa, which is essential for male fertility. Previous studies have shown that Na^+^,K^+^-ATPase α4 is highly expressed in the sperm flagellum; however, the spatial arrangement of Na^+^,K^+^-ATPase α4 at the subcellular level and its relationship to the functional state of the cells are unknown. We studied this here using stimulated emission depletion (STED) super-resolution microscopy. We show that, under non-capacitated conditions, Na^+^,K^+^-ATPase α4 is distributed in a trilinear pattern along the midpiece and as a scattered single line along the principal piece segment of the sperm flagellum. Under capacitated conditions, Na^+^,K^+^-ATPase α4 pattern undergoes remodelling and its distribution shifts into a single line along the entire length of the flagellum. On the other hand, Na^+^,K^+^-ATPase α1 the somatic isoform of Na^+^,K^+^-ATPase also present in sperm, exhibits a similar trilaminar localization at the flagellar midpiece but a bilinear pattern in the principal piece. This distribution, unlike that of Na^+^,K^+^-ATPase α4, does not change during sperm capacitation. These differences in the localization pattern and spatial dynamics of Na^+^,K^+^- ATPase isoform expression highlights the dissimilarities in the roles of both ion transporters. The specific modulation of Na^+^,K^+^-ATPase α4 distribution, combined with the unique role that it has in sperm function, stresses the importance of Na^+^,K^+^-ATPase α4 for male fertility.

**Significance statement:** This is the first demonstration of the highly structured nature of Na^+^,K^+^-ATPase in the plasma membrane of sperm, including the sperm specific Na^+^,K^+^-ATPase α4 isoform, which is key for male fertility, and the somatic Na^+^,K^+^-ATPase α1, which is present in all cells. Utilizing stimulated emission depletion (STED) super resolution microscopy, we discovered that Na^+^,K^+^- ATPase α4 and Na^+^,K^+^-ATPase α1 have different distributions along the sperm flagellum. Moreover, only Na^+^,K^+^-ATPase α4 undergoes remodelling during sperm capacitation. These specific patterns of localization that are dependent on the sperm functional state in combination with the different function and regulation of Na^+^,K^+^-ATPase isoforms highlights the sophisticated mechanisms that cells have evolved to fulfil their unique function.

## Introduction

Spermatozoa maintain a tight regulation of their intracellular ion composition, which is essential for their function and fertilization capacity. To achieve this, sperm express a series of ion transport proteins at their cell plasma membrane, some of which are unique and not shared with those of somatic cells (1–6). Among these transporters is Na^+^,K^+^-ATPase (EC/7.2.2.13), an integral plasma membrane protein complex that utilizes the energy from ATP hydrolysis to move 3 Na^+^ out of the cell in exchange for 2 K^+^ that are brought in (7–9). The asymmetrical Na^+^ and K^+^ distribution that Na^+^,K^+^-ATPase creates across the cell surface is crucial for various cellular and physiological processes including maintaining cell ion homeostasis, cell volume, resting membrane potential, and sodium-dependent secondary transport of different solutes and ions across the cell plasma membrane(10, 11).

The ion transport function and enzymatic hydrolysis of ATP catalyzed by Na^+^,K^+^-ATPase primarily depend on the α polypeptide or catalytic subunit of Na^+^,K^+^-ATPase. The other main subunit that constitutes Na^+^,K^+^-ATPase, the glycosylated β subunit, works as a chaperone molecule that facilitates the folding and delivery of the whole Na^+^,K^+^-ATPase transport complex to the plasma membrane (7, 12, 13). Mammalian cells express multiple isoforms of the Na^+^,K^+^- ATPase α (α1, α2, α3, and α4) and β (β1, β2, and β3) subunits. The pairing of Na^+^,K^+^-ATPase α and β isoforms in different combinations generates distinct Na^+^,K^+^-ATPase isozymes, each of which exhibits a tissue-specific and developmentally regulated pattern of expression and have distinct functional properties(14–17).

Mammalian male germ cells of the testis are the only cells that express the Na^+^,K^+^- ATPase α4 isoform. In addition, they also express the ubiquitous Na^+^,K^+^-ATPase α1 polypeptide of somatic cells and two of the β subunits (β1 and β3) (18). Na^+^,K^+^-ATPase α4 and α1 exhibit different affinities for Na^+^ and K^+^, with Na^+^,K^+^-ATPase α4 having a greater affinity for intracellular Na^+^ activation and a lower affinity for K^+^ activation compared to Na^+^,K^+^-ATPase α1 (7, 17). These particular biochemical properties better adapts Na^+^,K^+^-ATPase to the function of sperm. We have previously shown that genetic deletion of Na^+^,K^+^-ATPase α4 in mice results in complete infertility of male mice, with female mice being unaffected (18, 19). Moreover, sperm from Na^+^,K^+^-ATPase α4 knock-out mice are unable to fertilize oocyte in-vitro (18). The main consequence of Na^+^,K^+^-ATPase α4 disruption is a severe reduction of sperm total and hyperactive motility, a characteristic pattern of sperm movement that sperm acquires after capacitation, which is essential for fertilization (18, 20, 21). The mechanisms underlying this mouse model of asthenospermia are derived from the dissipation of the transmembrane Na^+^ gradient, depolarization of plasma membrane, cytoplasmic acidification, elevated Ca^2+^ levels, and reduced ATP production (22, 23).

While previous studies have explored the overall expression and physiological relevance of Na^+^,K^+^-ATPase α4 in sperm function, there is little information regarding the subcellular organization of this protein at the plasma membrane. Na^+^,K^+^-ATPase α4 is localized mainly in the midpiece of the mouse sperm flagellum; however, these studies were limited by the use of low-resolution microscopy (24, 25). To determine the spatio-functional relationship of Na^+^,K^+^- ATPase α4, we here examined the 3D distribution of Na^+^,K^+^-ATPase α4 within the sperm flagellum using stimulated emission depletion (STED) super-resolution microscopy. The nanometer-scale resolution that can be obtained with this equipment revealed that in non-capacitated sperm, Na^+^,K^+^-ATPase α4 shows a trilinear pattern along the flagellar midpiece and a single line at the principal piece of the sperm flagellum. Interestingly, this distribution is subjected to remodeling when sperm undergoes capacitation, with Na^+^,K^+^-ATPase α4 being reorganized in a single column along the entire length of the sperm flagellum. In contrast, Na^+^,K^+^-ATPase α1 has a trilinear distribution at the midpiece and bilinear localization at the principal piece, which is independent of the capacitated state of the cells.

## Results and Discussion

### Na^+^,K^+^-ATPase *α*4 is organized in three columns in the midpiece of non-capacitated *sperm*

Na^+^,K^+^-ATPase α4 has been shown to be expressed at the plasma membrane of the sperm flagellum, where it plays a critical role in sperm motility. However, previous studies were limited to the use of conventional fluorescence microscopy (24–26), which does not provide the resolution needed to distinguish compartmentalization of proteins within different regions of the cell plasma membrane. This resulted in images that identified Na^+^,K^+^-ATPase α4 evenly present along the midpiece of the sperm flagellum, with lower expression in the principal piece (25). In contrast, STED microscopy provides a resolution of 40 nm surpassing the diffraction-limitation of regular fluorescent microscopes, enabling the visualization of proteins with enhanced resolution, and allowing to establish their subcellular organization at the level of nanometer scale (27, 28). This is particularly important when considering cells like mouse spermatozoa, in which the flagellum has a width of ∼1 µm and a total length of ∼120 µm (29, 30).

To identify Na^+^,K^+^-ATPase α4, we here used an antibody that we made against a N-terminal sequence (RPSTRSSTTNRQPKMKRR) of Na^+^,K^+^-ATPase α4, which shows excellent Na^+^,K^+^-ATPase selectivity (*Fig. S1A*). To further determine the specificity of this antibody, we tested its reactivity toward sperm from a mouse which we had previously engineered to ablate the expression of Na^+^,K^+^-ATPase α4. Our results showed that the antibody showed no label in Na^+^,K^+^-ATPase α4 knock-out sperm but nicely identified the protein in wild type sperm. This showed that the anti-Na^+^,K^+^-ATPase α4 antibody has the required specificity in identifying our protein of interest and therefore, appeared amenable for use in STED microscopy (*Fig.S1B*).

Interestingly, our analysis of wild type sperm with STED microscopy revealed that, rather than being evenly distributed over the cell plasma membrane, Na^+^,K^+^-ATPase α4 exhibits a highly organized cell surface pattern of distribution (*movie S1*). Thus, in sperm collected from the caudal portion of mouse epididymis and maintained under non-capacitated conditions, Na^+^,K^+^-ATPase α4 is organized in three lanes that run along the midpiece section of the sperm flagellum (*Fig 1A,1B*). This trilinear organization becomes discontinuous entering the principal piece and appears at that point as a single line (*Fig 1C*).

**Figure 1:**
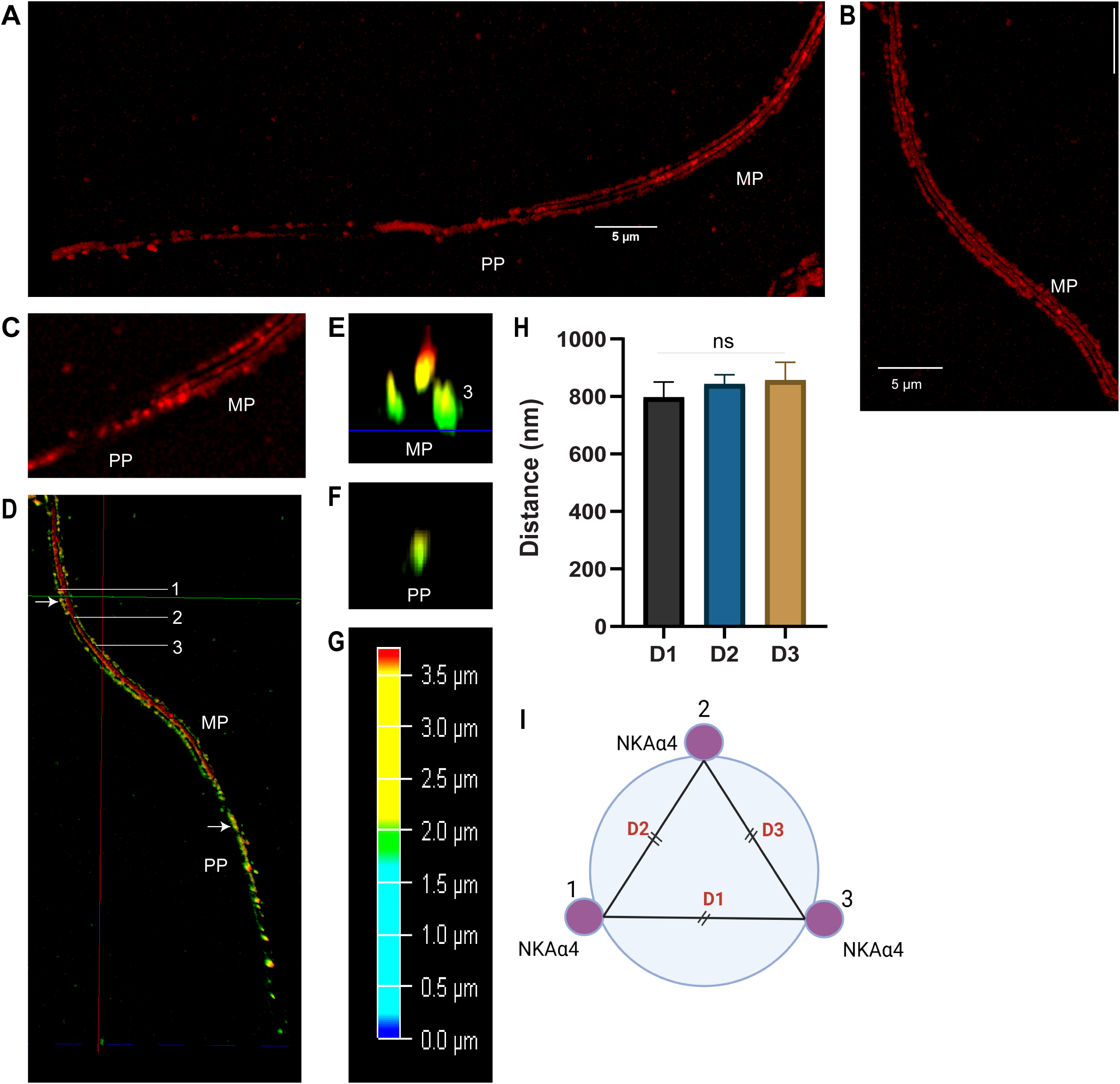
Na^+^,K^+^-ATPase α4 organization in sperm flagellum under non-capacitated condition. **(A)** 3D reconstructed image of the sperm flagellum showing distribution of Na^+^,K^+^- ATPase α4 (NKAα4) along the whole length of the flagellum, including the midpiece (MP) and principal piece (PP). **(B)** Trilinear distribution of Na^+^,K^+^-ATPase α4 in the midpiece (MP). The scale bars represent 5 µm. **(C)** Sperm flagellar segment showing the mid-principal piece boundary demonstrates the three lanes merging into a single lane at the end of midpiece. **(D)** The depth color coded 3D image of Na^+^,K^+^-ATPase α4 is showed in the whole flagellum. **(E)** The y-z cross section of the image obtained from D, sectioned at the midpiece (at the site marked with an arrow in D) showing three tight clusters (1, 2 and 3) of Na^+^,K^+^-ATPase α4. **(F)** The y-z cross section of the image obtained from the principal piece (PP) (at the site marked with an arrow in D) showing one cluster. Colors in D, E and F represent the z positions (refer to the color scale bar **G**). **(H)** Distance between Na^+^,K^+^-ATPase α4 clusters measured from the cross section of 3D images using LAS X analysis software. D1 represents the distance between lane 1 and 3, D2 is the distance between lane 1 and 2 and D3 is the distance between lane 2 and 3. Bars represent the mean ±SEM for n=8. ns indicates no significant statistical differences between values, with *P* < 0.05. **(I)** Schematic of the spatial distribution of three Na^+^,K^+^-ATPase α4 (NKAα4) clusters at the cross section of the midpiece shown in (E). The clusters or the lanes are equally distant from each other as derived from (H), forming an equilateral triangle connecting them as depicted in (I). The schematic was created with BioRender.com.

To determine the spatial position of the protein lanes, we obtained cross sections of the 3D STED images of sperm midpiece and principal piece. These images are shown as depth coded along the *z* axis direction (*Fig 1D*). The arrows in fig 1D mark the sites of the cross section from midpiece and principal piece resulting in fig 1E and 1F respectively. As shown, Na^+^,K^+^-ATPase α4 is arranged in three tight clusters in the midpiece that we designated as 1, 2 and 3 (*Fig 1E*) whereas as one compact cluster in the principal piece (*Fig 1F*). From the color coded of depth, it can be estimated that cluster 1 and 3 lie parallel to each other being at the same level of z axis, whereas cluster 2 is on the top of the z axis (*Fig 1E,1G*). The distances between the three lanes were determined using the Leica Application Suite X (LAS X) analysis tool. This showed that the distance between lanes 1 and 3 (shown as D1) was 797 nm, between lanes 1 and 2 (designated D2) was 843 nm, and between lanes 2 and 3 (named D3) was 856 nm (*Fig 1H*). A one-way anova test showed that the mean distances were not statistically different. Therefore the Na^+^,K^+^-ATPase α4 arrangement shows that the lines are equally spaced. Figure 1I illustrates a schematic of the three clusters in the longitudinal cross-section, where the distances between the lanes (1, 2, and 3) are represented as D1, D2, and D3, respectively, forming an equilateral triangle. For further confirmation, we also determined the angle between the lanes from the cross section using the LAS X software and found that the angle between the lanes is ∼60 ° suggesting the proteins are spatially organized in three equally spaced lanes.

The cross section of the mouse sperm flagellum is approximately circular (31). Based on the distribution of Na^+^,K^+^-ATPase α4, which is present at the sperm plasma membrane (32), the diameter of the mouse sperm flagellar midpiece can be predicted using the formula:

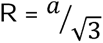

Where, *R* is the radius of the circle and *a* is the side of the equilateral triangle inscribed within the circle. Taking the length of the side 832 (mean of D1, D2 and D3), we calculated a radius of 480 nm and hence the diameter of the midpiece stands at approximately 960 nm. This supports previous finding where the width of the mouse sperm tail was reported to go from 900 nm to 1 µm in mice (30).

The distinct arrangement of Na^+^,K^+^-ATPase α4 shows the remarkable high organization that this ion transporter has in sperm. Previous studies have reported a particular arrangement for other ion transporters. For instance, the sperm-specific Ca^+^ channel CatSper exhibits a quadrilateral organization along the principal piece of the sperm flagellum (33). Likewise, the sperm H^+^ channel Hv1 displays an asymmetric bilateral distribution also in the flagellar principal piece. These patterns are uniform irrespective of the capacitation status of the cells (34).

### Na^+^,K^+^-ATPase ***α***4 is redistributed upon capacitation

We also determined the spatial organization of Na^+^,K^+^-ATPase α4 after incubating mouse sperm in medium which supports capacitation. To achieve capacitation, sperm was incubated with Card FertiUp Preincubation Medium, which provides optimal conditions and is successfully used to capacitate sperm for *in vitro* fertilization purposes (35, 36). Interestingly, under capacitated conditions, we found dramatic changes in the pattern of Na^+^,K^+^-ATPase α4. The trilinear arrangement observed under the non-capacitation state, now reorganized as a single line along the entire length of the sperm flagellum with predominant presence in the midpiece (*Fig 2A,2B*). The cross section at the midpiece and principal piece from the depth coded 3D reconstructed image (*Fig 2B*) showed one compact cluster (*Fig 2C*). To confirm the capacitated status of the cells, we evaluated their capacity to develop hyperactive motility, a characteristic high amplitude-low frequency movement that sperm acquires at capacitation (21, 22, 37). Figure 2E shows that sperm acquired hyperactivated motility as measured using computer assisted sperm analysis (CASA). This confirmed the capacitated state of the cells under our study conditions.

**Figure 2:**
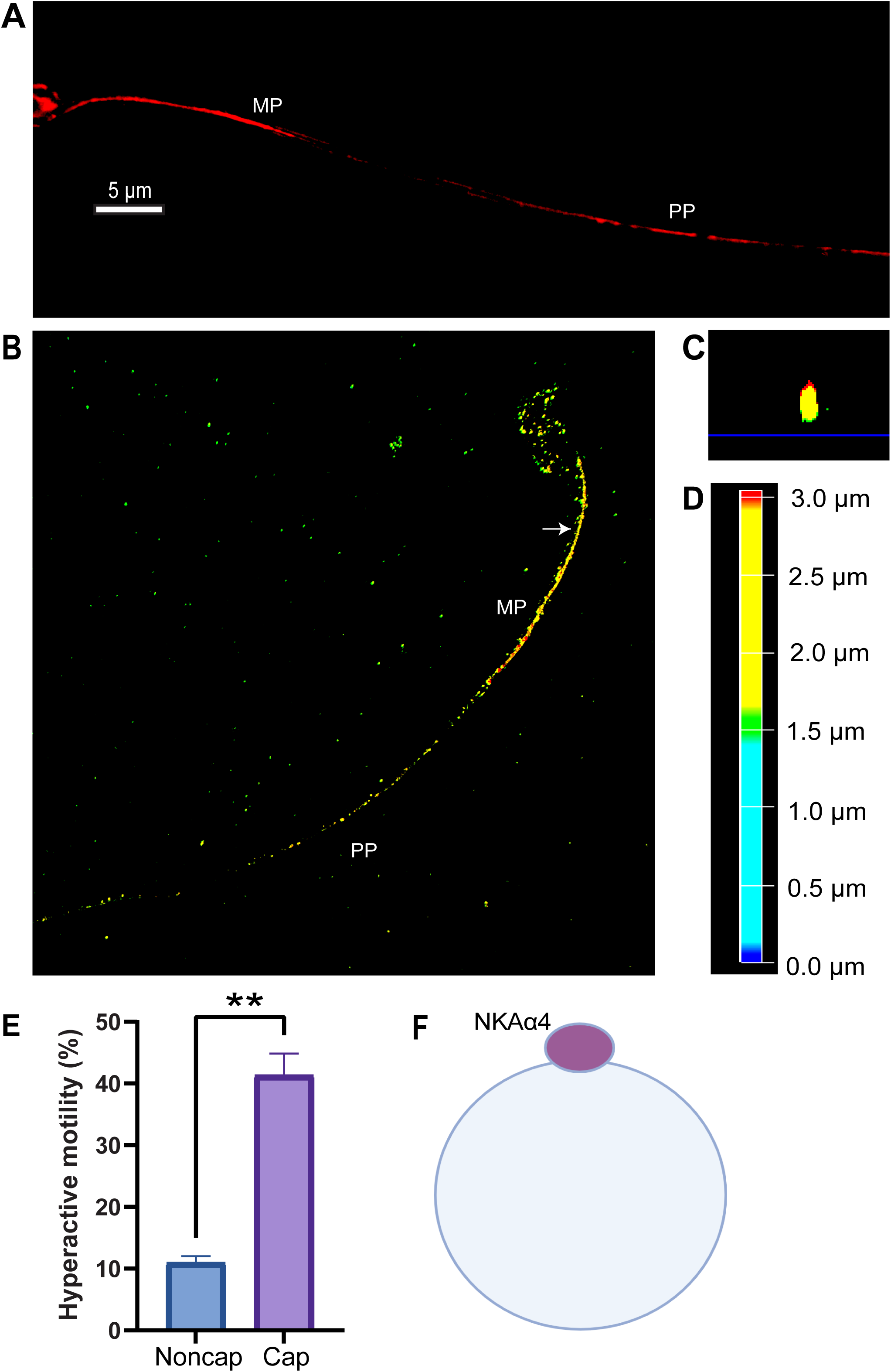
Na^+^,K^+^-ATPase α4 is redistributed upon sperm capacitation. **(A)** 3D reconstructed image of Na^+^,K^+^-ATPase α4 along the entire flagellum in capacitated sperm. **(B)** Depth coded 3D image of Na^+^,K^+^-ATPase α4. **(C)** The y-z cross section of the flagellum showing a compact Na^+^,K^+^-ATPase α4 cluster. The site of cross section is marked with an arrow in B. The scale bars are 5 µm. Colors in B and C represent the z positions (see color scale bar **D**). **(E)** Hyperactive motility in non-capacitated (Noncap) and capacitated (Cap) sperm evaluated with CASA. **(F)** A schematic showing the three lanes to be remodeled into one. The schematic was created with BioRender.com.

The capacitation dependent redistribution of other proteins, such as sperm surface antigens has been reported in boar, pig, hamster, mouse, and rat (38–41). For instance, PT-1, a protein restricted to the last segment of the sperm tail appears in the entire flagellum after capacitation (42). Cytoskeletal proteins, such as actin undergo dynamic remodeling during capacitation (43). Regarding Na^+^,K^+^-ATPase α4, we have previously demonstrated that the ATPase activity of this isoform is up-regulated at capacitation and that this is accompanied by an increase in expression of the transporter at the cell plasma membrane (32). Since sperm are terminally differentiated, transcriptionally inactive cells, the increase in Na^+^,K^+^-ATPase α4 activity will depend on capacitation associated changes that result from a shift of Na^+^,K^+^-ATPase α4 to specific areas of the plasma membrane during capacitation (32, 44). Our data here directly demonstrates the redistribution of Na^+^,K^+^-ATPase α4 when sperm capacitates (schematically shown in *Fig 2F*). The rearrangement of Na^+^,K^+^-ATPase α4 may provide an advantage in enhancing the transmembrane Na^+^ gradient at specific microdomains of the sperm cytosol, which will provide the chemical force that fuels the activity of other sperm ion transport systems, such as the Na^+^/H^+^ and Na^+^/Ca^2+^ exchanger (45–47). These ion transporters are essential in increasing the sperm cytosol pH and Ca^2+^ levels that are required to achieve the capacitated state. In addition, the K^+^ gradient that Na^+^,K^+^-ATPase α4 generates assists K^+^-channels, such as Slo3, which is critical for the hyperpolarization that cells acquire at capacitation (48, 49).

The most prominent changes that we found for Na^+^,K^+^-ATPase α4 corresponds to its expression as a single line in the flagellar principal piece. Structurally, the principal piece of the flagellum is more flexible than the midpiece segment (50, 51). This allows this flagellar region to be primarily involved in hyperactive motility during capacitation. Coincidentally, ion transporter molecules such as NHE, Hv1, Slo3 or CatSper, which are directly involved in sperm hyperactive motility are also found in the flagellar midpiece (33, 34, 49, 52, 53). Interestingly, the changes in the distribution pattern of Na^+^,K^+^-ATPase α4 have not been reported for Hv1 or CatSper (33, 34). Therefore, it appears that capacitation dependent changes in protein distribution is a phenomenon limited to selected sperm proteins like Na^+^,K^+^-ATPase α4, which is crucial for sperm hyperactivation and the sperm capacity to fertilize the oocyte.

### Na^+^,K^+^-ATPase ***α***1 also has a distinct pattern of expression in the sperm flagellum independent of the capacitated status of the cells

To assess the pattern of expression of the other Na^+^,K^+^-ATPase α isoform expressed in sperm, we investigated the distribution of Na^+^,K^+^-ATPase α1 along the sperm flagellum under non-capacitated and capacitated conditions. Our results show that Na^+^,K^+^-ATPase α1 is present along the entire length of the sperm flagellum as has been previously described (26). Like Na^+^,K^+^-ATPase α4, Na^+^,K^+^-ATPase α1 is also distributed as three lines in the midpiece of the sperm (*Fig 3A,3B*). The cross section of the depth coded images of the flagellar midpiece revealed three compact dots (denoted by 1, 2 and 3) showing the trilinear organization (*Fig 3C,3D*). From the depth coded z axis, it is found that lanes 1 and 3 are at the same depth level whereas Lane 2 is on the top of the axis (*Fig 3D and 3F*). Interestingly, beyond the flagellar midpiece and in the principal piece, Na^+^,K^+^-ATPase α1 follows a bilinear distribution which appears to be spatially at the same level (*Fig 3E,3F*).

**Figure 3:**
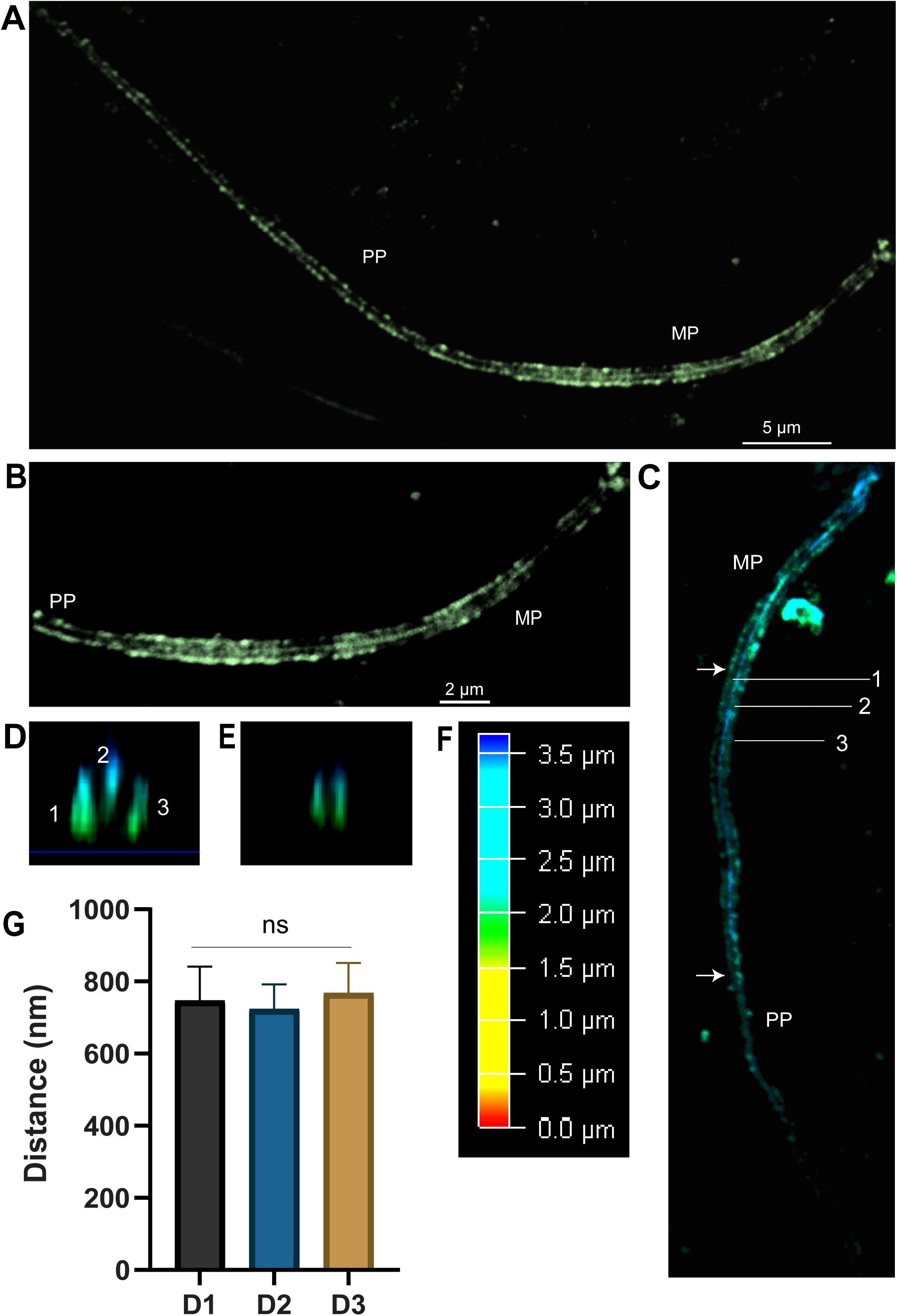
Na^+^,K^+^-ATPase α1 has a different organization at the sperm flagellum than Na^+^,K^+^-ATPase α4 under non-capacitated condition. **(A)** 3D distribution of Na^+^,K^+^-ATPase α1 along the entire flagellum. **(B)** Na^+^,K^+^-ATPase α1 localization in the midpiece (MP) showing a trilinear pattern in the midpiece (MP) and a bilinear pattern at the principal piece (PP). Scale bar represents 5 µm (A) and 2 µm (B) respectively. **(C)** Color coded 3D image showing the position of Na^+^,K^+^-ATPase α1 along the z direction (see color bar in **F** for reference). **(D)** y-z cross section obtained from (C) shows three domains (1,2 and 3) at the midpiece (MP). **(E)** Cross section showing Na^+^,K^+^-ATPase α1 distributed as two lines at the principal piece of the flagellum. Arrows in C denote the sites of cross sections that result in D and E. **(G)** Distance between the lanes or clusters measured from the cross section of 3D images using LAS X analysis software. D1 represents the distance between lane 1 and 3, D2 is the distance between lane 1 and 2 and D3 is the distance between lane 2 and 3. Values are expressed as mean ±SEM for n=8. ns shows no significant statistical differences, with *P*L<L0.05.

Like Na^+^,K^+^-ATPase α4, the distances between the lanes for Na^+^,K^+^-ATPase α1, measured with LASX software, were found to be statistically similar, indicating that the three clusters are equally spaced from each other (*Fig 3G*). However, unlike Na^+^,K^+^-ATPase α4, the localization pattern of Na^+^,K^+^-ATPase α1 was preserved upon sperm capacitation (*Fig 4A-E*). In addition, we found that Na^+^,K^+^-ATPase α1 distribution remains unchanged when we repeated the experiments in sperm from the Na^+^,K^+^-ATPase α4 knock out mouse (*Fig S1C-E*). This indicates that the distribution of Na^+^,K^+^-ATPase α1 and α4 is not interdependent.

**Figure 4:**
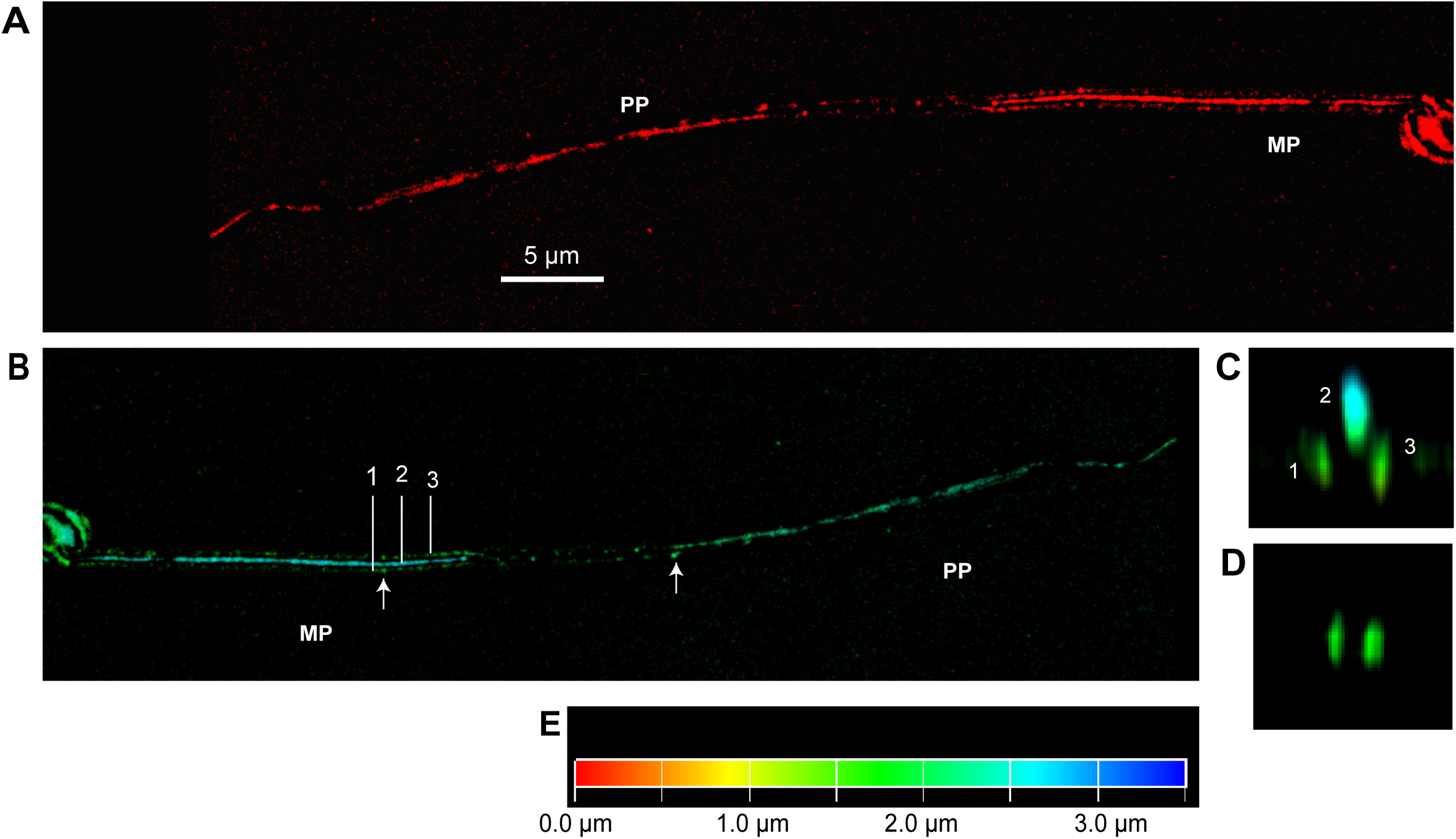
The pattern of distribution of Na^+^,K^+^-ATPase α1 is not changed upon sperm capacitation. **(A)** 3D organization of Na^+^,K^+^-ATPase α1 along the sperm flagellum, showing a trilinear pattern in the midpiece which is discontinued into two lanes in the principal piece. Scale bar represents 5 µm. **(B)** Expression of Na^+^,K^+^-ATPase α1 color-coded along the z axis (refer to color bar **E** for depth of field). **(C)** y-z cross section at the midpiece showing three domains (1, 2 and 3) **(D)** y-z cross section at the principal piece showing only two Na^+^,K^+^-ATPase α1 clusters. Arrows in B shows the sites of cross section from midpiece (C) and principal piece (D) respectively.

The difference between Na^+^,K^+^-ATPase α1 and Na^+^,K^+^-ATPase α4 with respect to sperm capacitation is interesting. Earlier results from our laboratory have shown that, while Na^+^,K^+^- ATPase α1 maintains the basal ion concentrations in the cell, Na^+^,K^+^-ATPase α4 is dedicated to sperm motility, hypermotility and capacitation(19). Therefore, the repositioning of Na^+^,K^+^-ATPase α4 in the sperm flagellum during capacitation, may be related to its sperm specific roles and the need to maintain a steeper Na^+^ gradient in specific intracellular domains, which is required to prepare sperm to fertilize the egg.

## Conclusion

We here report that both of the Na^+^,K^+^-ATPase α isoforms expressed in sperm have a different spatial and functional pattern of distribution along the flagellum. While both share a trilinear organization in the midpiece of the flagellum, they differ in their localization at the flagellar principal piece. Moreover, Na^+^,K^+^-ATPase α4 but not Na^+^,K^+^-ATPase α1 is remodeled during sperm capacitation, to adopt a distribution as a single line along the entire length of the sperm flagellum. This particular spatial distribution of Na^+^,K^+^-ATPase α4 and its redistribution upon sperm capacitation may respond to the critical physiological role that this isoform plays in sperm movement and fertility.

## Materials and Methods

### Sperm preparation and invitro capacitation

Sperm from the caudal portion of the epididymis were isolated from adult wild type and Na^+^,K^+^-ATPase α4-KO male mice (12-14 weeks old, C57Bl/6 J) as previously reported(54). Briefly, sperm were collected by swim-up in the modified Whitten’s Media containing 100mM NaCl, 4.7mM KCl, 1.2mM KH_2_PO4, 1.2mM MgSO4, 5.5mM glucose, 0.8mM pyruvic acid, 4.8mM lactic acid, 20mM Hepes, and 1.7mM CaCl_2_. To induce capacitation, cells were incubated in CARD FERTIUP Preincubation Medium: PM (Cosmo Bio LTD, Koto-ku, Tokyo, Japan) at 37°C for 10 mins and then incubated at the same temperature in 5% CO_2_ for an hour.

### Immunocytochemistry

For Immunolabelling, sperm was washed in Phosphate Buffered Saline (PBS) (ThermoFisher Scientific, Waltham MA, USA) twice (120x g, 5 mins for each wash) at 4°C and fixed in 4% paraformaldehyde (PFA) for 15 min. The cells were then washed three times in PBS (380x g, 5mins for each wash) and then resuspended in PBS. The diluted cells in suspension were placed on a slide and air-dried. To minimize autofluorescence the slides were exposed to light emitting diode (LED) light for 48 hrs. Then, samples were permeabilized with 0.3% Triton-X for 10 min in PBS, washed three times with PBS, and then blocked with 2% BSA in PBS for 1h at RT. The cells were incubated at 4°C overnight in a humid chamber with the primary antibodies. These included anti-ATP1A4 (in-house antibody raised against epitope RPSTRSSTTNRQPKMKRR of Na^+^,K^+^-ATPase α4, used at a 1:1500 dilution); and anti-ATP1A1 (ProteinTech, Rosemont IL, USA; dilution of 1:1500). The samples were then washed three times with PBS and incubated at RT for 1h with Goat Anti-Rabbit Alexa Flour 594 conjugated secondary antibody (Invitrogen, Carlsbad CA, USA) diluted to 1:1000 in PBS with 2% BSA. Cells were then washed three times in PBS, and then mounted with Prolong Gold (ThermoFisher, Waltham MA, USA), and cured for 24 hours for imaging with STED microscopy. For Nikon Eclipse 80i Fluorescence Microscope, the sample was mounted with Dapi and kept at 4°C and imaging was done at 20x magnification.

### Immunoblot

To determine the selectivity and specificity of in-house antibody, protein expression of Na^+^,K^+^-ATPase α4 in different tissue samples was evaluated by SDS-PAGE and immunoblotting as described previously(54). Briefly, tissues from kidney, liver, heart, brain, testis and sperm from wild type mice are homogenized and treated with lysis buffer. Samples were run on 8% SDS-PAGE gel and transferred on a PVDF membrane. Blocking for an hour in 5% blotto with TBST followed by overnight incubation with primary anti-ATP1A4 antibody at 4°C were performed. Horseradish peroxide conjugated anti-goat secondary antibody (Jackson ImmunoResearch Europe Ltd, Cambridgeshire, UK) at 1:2000 dilution and chemiluminescence were used for detection.

### Sperm-motility assay

Non-capacitated or capacitated sperm from wild type mice were analyzed for sperm motility using Computer Aided Sperm Analysis tool or CASA (version 3.9.8, Penetrating Innovations) as previously described (25). Briefly, 1X10^6^ spem were labelled with 2 μl of a 75-μM stock of SITO 21, a nucleic acid staining fluorescence to track sperm movement. After incubating for 2 mins with the dye the sample is visualized under microscope and sperm hyperactive motility is then analyzed with the CASA.

### STED Microscopy and image analysis

One-color STED images were recorded with a Leica TCS SP8 STED 3X Super Resolution Microscope with an oil immersion objective (HC PL APO 100x/1.40 OIL CS2 objective, Oil immersion type F; Leica Wetzlar, Germany). The bit depth was set to 12. To get sharp and more sensitive images, the HyD was turned on as the photodetector. The HyD (Hybrid Detector) provides high contrast image with low-dark noise and reduce photobleaching of the sample. The White Light Laser (WLL) had an excitation wavelength of 598±4 nm for Alexa Flour 594 and a STED Laser Beam of 775 nm was used for depletion without gating. A pixel size of 170 to 180 nm was used with dwell time of 3.16 µs and each line was scanned for 3 times (Line Accumulation). The pinhole was set to 1.0 AU. The z-step size was 0.16 µm. XY format was optimized and set as suggested by STED. Speed was set to 400 Hz, meaning repeated scanning with depletion laser for 400 times per second which facilitates faster imaging. The image is then deconvoluted which reveals the fine structure of the image reducing the background noises which might be otherwise missed. For 3D visualization and depth coding Leica Image Analysis software was used. For determining the distance, and angle between the three lines, we have used LASX Leica Analysis Software and for other image processing an open software, Fiji ImageJ was employed.

### Statistical Analysis

Experiments were repeated at least three times and the statistical significance of mean and differences between distances were determined by one-way anova test. For determining the significance of difference between hyperactive motility, paired t-test using GraphPad Prism (San Diego, CA, USA) was employed. Statistical significance was defined as *p* < 0.05.

## Acknowledgement

We thank Dr. Sarah Tague, Dr. Christine Smoyer and the Imaging Core of the Kansas Intellectual and Developmental Disabilities Research Center (KIDDRC) for their support. This core was supported by NIH grant S10 OD023625 to the University of Kansas Medical Center, Kansas City.

