## Supplementary figure 1 for "The sperm specific Na^+^,K^+^-ATPase α4 shows a highly structured and dynamic distribution at the sperm flagellum"

Mumtarin J Oishee, Jeffrey P. McDermott, Gladis Sanchez & Gustavo Blanco.
Department of Cell Biology and Physiology, University of Kansas Medical Center, Kansas City, KS 66160, USA.

**Correspondence to**: Gustavo Blanco, Department of Cell Biology and Physiology, 3901 Rainbow Boulevard, Kansas City, Kansas 66160.

**This PDF file includes:**
Figure S1
Legend for Movie S1

**Other supporting materials for this manuscript include the following:**
Movies S1

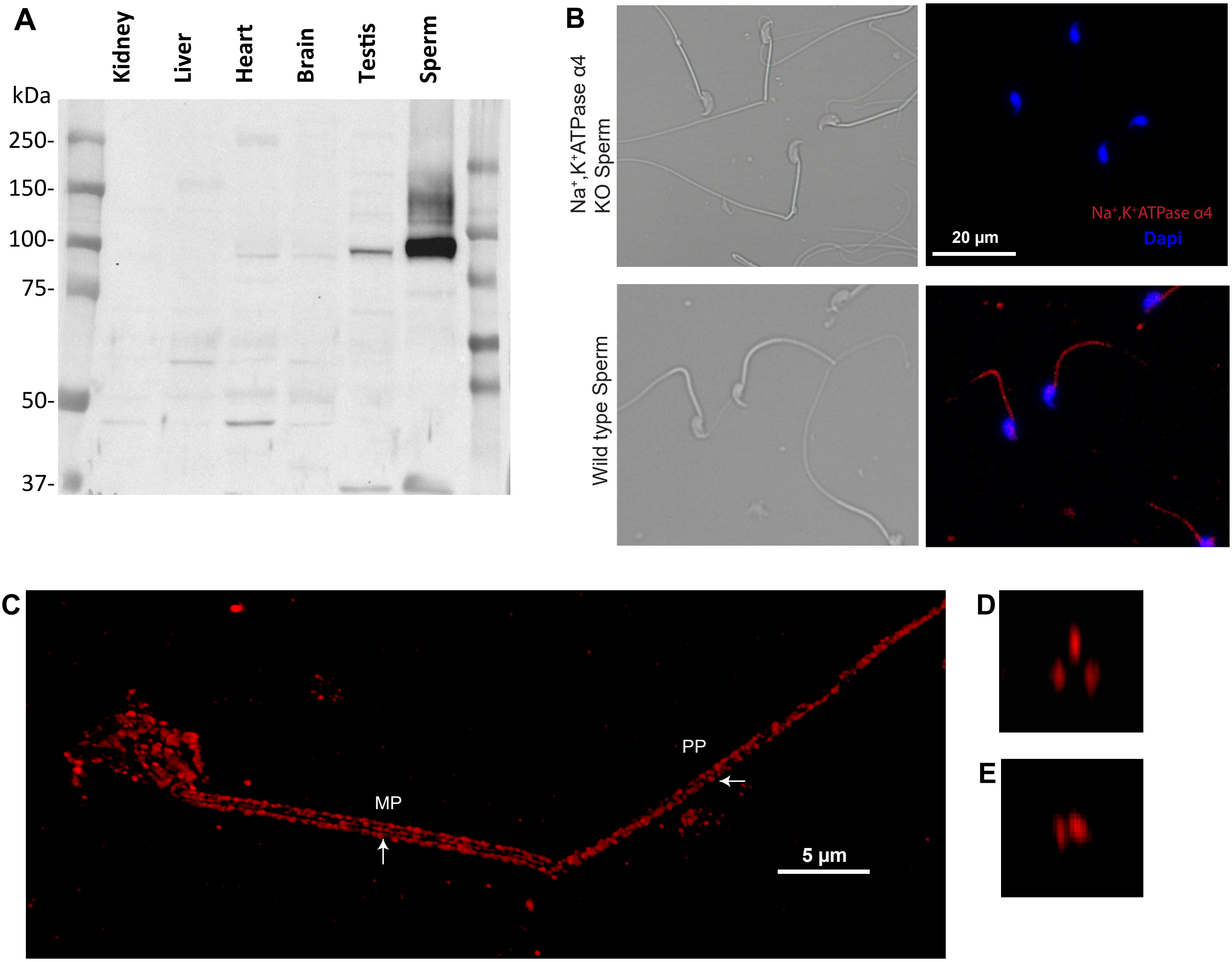


**Supplementary Fig S1: Expression of Na$^+$,K$^+$-ATPase in wild type and Na$^+$,K$^+$-ATPase α4 Knock out sperm**. **(A)** Immunoblot showing the expression of Na$^+$,K$^+$-ATPase α4 (≈ 100KDa) only in sperm and testes. Protein was isolated from different tissue samples and run on a 8% SDS/ polyacrylamide gel. Membranes were probed with anti-Na$^+$,K$^+$-ATPase α4 antibody raised in rabbit. Chemiluminescence assay using HRP-conjugated secondary antibody was used for detection. **(B)** Sperm from the KO and wild type mice were probed with anti- Na$^+$,K$^+$-ATPase α4 antibody followed by a goat-anti rabbit Alexa 594 conjugated secondary antibody and co-stained with DAPI at 20X magnification using Nikon Eclipse 80i Fluorescence Microscope. Scale bar represents 20 µm. **(C)** 3D image of Na$^+$,K$^+$-ATPase α1 expressed in the KO sperm shows trilinear pattern in the midpiece and a bilinear pattern in the principal piece, which is confirmed from the cross section at the midpiece **(D)** and principal piece **(E)** respectively. The sites for cross section are shown in C with arrow. Scale bar represents 5 µm.

**Movie S1 (separate file)**

This movie demonstrates the 3D distribution of $Na^+$,$K^+$-ATPase α4 (in grey) in two sperm. Sperm midpiece exhibits trilinear organization of this protein and the sperm principal piece shows singular column but scattered distribution of $Na^+$,$K^+$-ATPase α4.
